# The hidden pangenome: comparative genomics reveals pervasive diversity in symbiotic and free-living sulfur-oxidizing bacteria

**DOI:** 10.1101/2020.12.11.421487

**Authors:** Rebecca Ansorge, Stefano Romano, Lizbeth Sayavedra, Maxim Rubin-Blum, Harald Gruber-Vodicka, Stefano Scilipoti, Massimiliano Molari, Nicole Dubilier, Jillian Petersen

**Affiliations:** Max Planck Institute for Marine Microbiology, Bremen, Germany; Division for Microbiology and Ecosystem Science, University of Vienna, Austria; Quadram Institute Bioscience, Norwich Research Park, Norwich, United Kingdom; Israel Limnology and Oceanography Research, Tel Shikmona, Haifa, Israel; Department of Bioscience, Aarhus University, Aarhus, Denmark; MARUM, University of Bremen, Bremen, Germany

## Abstract

Sulfur-oxidizing *Thioglobaceae*, often referred to as SUP05 and Arctic96BD clades, are widespread and common to hydrothermal vents and oxygen minimum zones. They impact global biogeochemical cycles and exhibit a variety of host-associated and free-living lifestyles. The evolutionary driving forces that led to the versatility, adoption of multiple lifestyles and global success of this family are largely unknown. Here, we perform an in-depth comparative genomic analysis using all available and newly generated *Thioglobaceae* genomes. Gene content variation was common, throughout taxonomic ranks and lifestyles. We uncovered a pool of variable genes within most *Thioglobaceae* populations in single environmental samples and we referred to this as the ‘hidden pangenome’. The ‘hidden pangenome’ is often overlooked in comparative genomic studies and our results indicate a much higher intra-specific diversity within environmental bacterial populations than previously thought. Our results show that core-community functions are different from species core genomes suggesting that core functions across populations are divided among the intra-specific members within a population. Defense mechanisms against foreign DNA and phages were enriched in symbiotic lineages, indicating an increased exchange of genetic material in symbioses. Our study suggests that genomic plasticity and frequent exchange of genetic material drives the global success of this family by increasing its evolvability in a heterogeneous environment.

## Introduction

Widespread high-throughput genome sequencing, metagenomics and comparative genome analyses have vastly increased our understanding of microbial biodiversity, revealing a complex landscape of genomic variation even among members of single species [1]. The collective gene content of lineages within a taxonomic group (e.g., a species) is commonly referred to as ‘pangenome’. This is divided into the ‘core genome’, defined as the set of genes encoded by all members of the group and the ‘accessory genome’, referring to the variable part that is only encoded in some members of the group. The mechanisms that lead to the emergence and evolution of accessory genomes and their role in the biology of species are the subject of ongoing debate [2], which gravitates around data obtained mainly from cultivated bacteria of medical relevance (e.g., *Escherichia coli*, *Mycobacterium tuberculosis*) [1, 3]. However, conclusions based on the study of isolates are not necessarily extendable to microbes in nature, as they neglect the diversity within populations, which is needed to understand a species’ evolutionary history [4]. To obtain a more realistic, comprehensive, and general view on the evolutionary driving forces that shape bacterial genome diversity, both free-living and host-associated populations need to be studied in their natural context [5].

Accumulating evidence suggests that there can be substantial subspecies diversity affecting function within environmental bacterial populations and microbiomes [6–8]. For example, populations of intracellular sulfur-oxidizing symbionts of bathymodiolin mussels harbor extensive gene content diversity within single host individuals [9–11]. These and other observations suggest that subspecies variation in symbiotic populations may be far more widespread than currently known [9]. These sulfur-oxidizing bacteria belong to a highly successful gammaproteobacterial group of symbiotic and free-living sulfur oxidizers widespread in the world’s oceans [12]. Based on the Genome Taxonomy Database [13], these bacteria belong to two well-known sister clades SUP05 and Arctic96BD within the *Thioglobaceae* family. Free-living *Thioglobaceae* are particularly abundant and active in oxygen minimum zones, anoxic marine zones and at hydrothermal vents playing a pivotal role in sulfur and nitrogen cycles [14–18]. To date, only three strains from the *Thioglobaceae* family have been isolated [19–21]. However, metagenomic and single-cell sequencing efforts revealed extensive metabolic versatility among free-living *Thioglobaceae* lineages, in particular sulfur, nitrogen, oxygen and carbon metabolism [22, 15, 16, 23, 24]. Underlining their versatility, members of this clade evolved symbioses with various deep-sea invertebrates which vary in their intimacy and life history, from vertical to horizontal transmission [25–27].

The metabolic diversity among free-living lineages [22, 15, 16, 24], the intraspecific gene content variation within the mussel symbionts [9–11], and the range of different lifestyles observed within the *Thioglobaceae* family indicate an impressive versatility and genomic plasticity. However, to date, a comprehensive understanding of this variability is lacking, as most studies focus on either symbiotic lineages or the free-living. Here we investigate the extent of genomic diversity and its relationship to bacterial lifestyles. We retrieved an extensive collection of *Thioglobaceae* genomes from public data and generated new metagenome-assembled genomes (MAGs) to assess the pangenome of this cosmopolitan marine microbial family. Specifically, we used high-resolution metagenomics, to determine the extent of genetic variation within and among *Thioglobaceae* sub-clades of different relatedness. We uncovered an unexplored sub-species diversity and interpret this in light of current evolutionary theories.

## Methods

### Sample collection and DNA extraction

Samples and collection sites are listed in the supplementary **Tab. S1**. An overview of the sampling locations is plotted as a geographic map using the tmap package (v. 2.0) in R (v 3.6.1) [28] (**Fig. 1**). We include newly generated sequencing data, publicly available metagenomes that were assembled and binned within our study and publicly available MAGs and genomes (details in **Tab. S1**). For newly generated sequencing data, mussels were collected, dissected on board and gill tissue was frozen and stored at −80°C or preserved in RNAlater and subsequently frozen at −80°C. DNA extraction was performed with the DNeasy Blood & Tissue kit (Qiagen, Germany) or according to Zhou et al. [29]. Library preparation and paired-end Illumina sequencing was performed at the Max Planck Genome Center Cologne. Sample acquisition and DNA extraction from mussel and sponge samples deriving from other studies are described in the respective publications [9, 10, 30–32] and summarized in **Tab. S1** and **S2**. To obtain published genomes we scanned publications and public databases (see **Tab. S3**). Accession numbers of published genome sequences included in this study are listed in **Tab. S2**.

**Figure 1.**
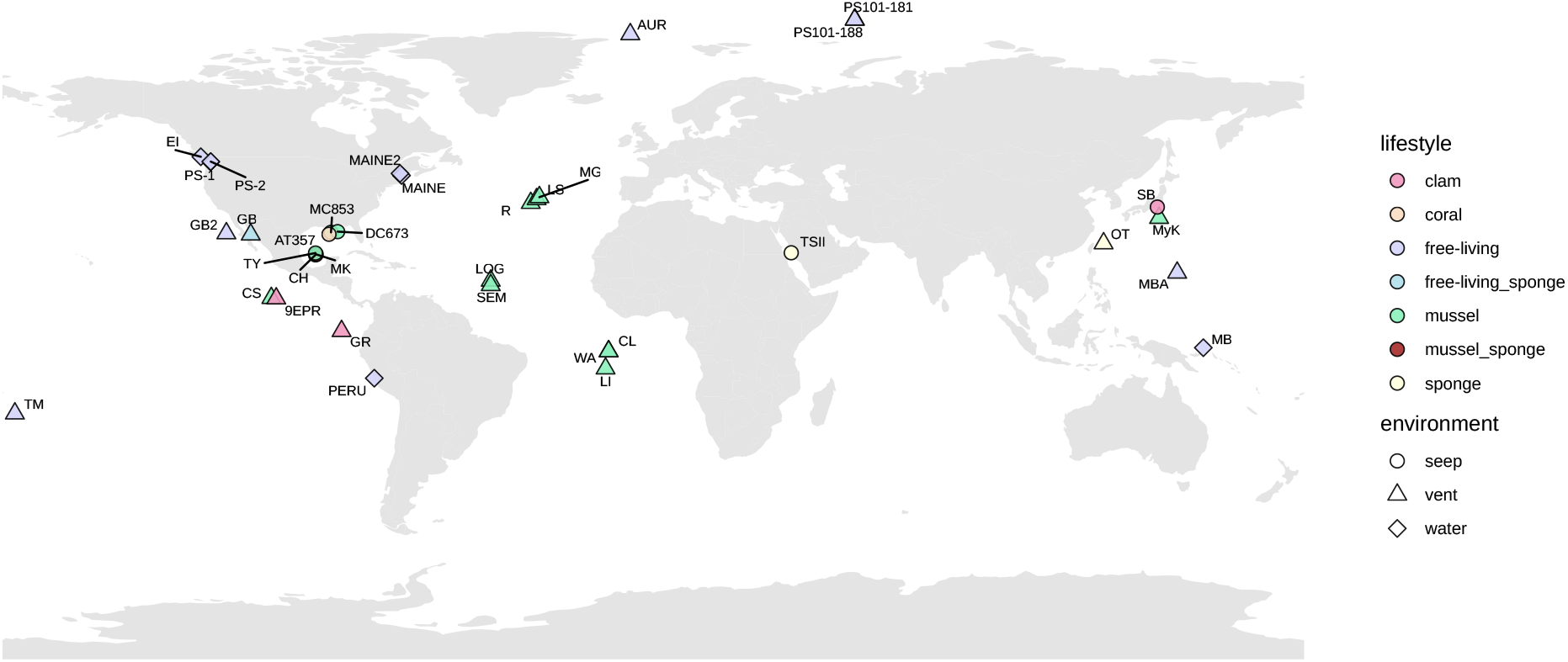
Sampling locations of all datasets included in the study. Color corresponds to lifestyle and shape of the symbols corresponds to the environment type. The names of the sites, their coordinates and additional metadata can be found in Tab. S1, S2, and S5.

### Metagenomics and genome binning

Libraries of genomic DNA were generated for each sample with the Illumina TruSeq DNA Sample Prep Kit (BioLABS, Germany). Details of the sequencing are shown in **Tab. S1**. Raw sequences were processed, assembled and binned either as described by the respective publication or as described in the following. For the metagenomes in this study, read adapters were removed and read quality was filtered to a minimum of two (Q2), using BBDuk (v 38.34, Bushnell B. - sourceforge.net/projects/bbmap/). Metagenomes were assembled for each sample individually with metaSPAdes using the default parameters and kmer sizes of 33, 55, 77, 99 and 127 [33, 34]. Binning of the sulfur-oxidizing symbiont MAGs in mussel hosts was performed using Bandage [35], differential coverage analysis combined with taxonomy and GC content [36] using gbtools [37], or Metabat2 [38]. This produced high-quality MAGs with the completeness of > 89% according to gammaproteobacterial marker genes in checkM (v 1.0.7) [39]. Publicly accessible genomes and MAGs included in this study were > 87% complete. We also included genome sequences from three vertically transmitted vesicomyid symbionts, which had lower checkM-estimated completeness (85-87%) reflecting their previously reported genome reduction [40, 41]. Genome contamination was corrected for the degree of strain diversity, as estimated by CheckM, and only genomes with a value < 5% were included in this study. Genome statistics of all MAGs used in this study are listed in **Tab. S1** and **S2**. Contigs smaller than 500 bp were excluded from all final MAGs created in this study.

### Pangenome analysis

For annotation consistency, all genomes, MAGs and SAGs were annotated with RAST-tk [43] as implemented in PATRIC [44]. Average nucleotide identity (ANI) and average amino acid identity (AAI) were calculated with the enveomics collection [45], and clusters of *Thioglobaceae* species were defined according to ANI cutoff > 95% [42, 43]. Percentages of conserved proteins (POCP) were calculated as described previously [44] for all genome pairs using a published bash script [45]. Amino acid sequences of all coding sequences were used to calculate core, accessory and unique genes among all genomes and MAGs within a single species cluster with BPGA (v 1.3.0) [46] using default settings of 0.5 sequence identity for clustering with USEARCH (v 1.1.1) [47]. To reflect the inability of assemblers to resolve microdiversity within an environmental sample and to separate these terms from the strain-specific gene analysis (see below) we referred to these sets as ‘assembly-based core’, ‘assembly-based accessory’ and ‘assembly-based unique’ genomes.

To obtain a representative set of genes encompassing all genomes of each species, we produced a single representative gene catalog per *Thioglobaceae* species, which included one protein sequence for each gene cluster observed in at least one genome. On these species-specific pangenome catalogs, we performed clustering of orthologous protein sequences to determine those shared between species using BPGA as described above. The resulting presence-absence matrix of gene clusters was formatted and shared genes among *Thioglobaceae* groups of interest were visualized, using the package UpsetR (v1.3.3) in R. The gene sets shared among all genomes of that species (species-specific gene catalogue) were uploaded to BlastKOALA (v2.2) [48] to assess the pathway modules and potential phenotypic discreteness of each proposed species (**Tab. S4**).

### Functional analysis

We mapped the genes of each MAG to the Kyoto encyclopedia of genes and genomes (KEGG) [49–51] using BPGA to assess and compare their functional genetic potentials. We performed two distance-based redundancy analyses (dbRDA) on Bray-Curtis dissimilarities of KEGG ortholog (KO) relative frequencies per MAG or genome, using the capscale function in the vegan (v2.5-6) package and the ggplot2 (v3.3.0) package in R. One dbRDA was performed on the KO profiles of all genomes in this study (*Thioglobaceae* family) and the second dbRDA was performed on the genomes of the SUP05 clade only (*Ca.* Thiomultimodus gen. nov.).

Permutational multivariate analysis of variance (PERMANOVA; 999 permutations) was performed on the Bray-Curtis dissimilarities of KO profiles in the SUP05 clade to determine the influence of ‘species’, ‘lifestyle’, and ‘environment’ (data in **Tab. S5**) on the genomic functional profiles, using the adonis2 function of the vegan package in R (**Tab. S6**). First, we performed a by ‘terms’ analysis. We observed that influence of ‘lifestyle’ on the data could be explained by the factor ‘species’ alone (**Tab. S6B**). This is because the factor ‘lifestyle’ was confounded by the factor ‘species’. Therefore, we excluded ‘lifestyle’ from the model and tested only the marginal effects of ‘species’ and ‘environment’ by performing a second PERMANOVA model by ‘margin’ (**Tab. S6C**). To visualize the effects of both factors on data clustering, we performed two additional dbRDA analyses - one conditioned for ‘species’ and the other one conditioned for ‘environment’, using the capscale function in the vegan package.

A generalized odds ratio for two-group ordinal data was computed using the R package genodds (v 1.0.0) on the functional KEGG KO frequencies to detect significant differences (Benjamini-Hochberg adjusted p-value < 0.05) between both lifestyles, symbiotic and free-living (**Tab. S7**).

### Phylogenetic tree calculation

We identified 98 core genes in a pangenome analysis, which included all genomes, MAGs, and SAGs in this study, supplemented with two outgroup genomes *Thiomicrospira arctica* (GCA_000381085.1) and *Thiomicrospira crunogena* (GCA_000012605.1). The protein sequences were aligned with MAFFT (v7.215) [52, 53] and sites of low confidence were identified and removed using GUIDANCE2 (v 2.02) [54]. The maximum likelihood tree was calculated using IQ-TREE (v1.6.9) [55] based on the concatenated protein sequences of the core genes, following the selection of the best evolution model by the ModelFinder module of IQ-TREE, using partition models [56] and ultrafast bootstrap [57].

### Protein domain prediction

We used Interproscan (v 5.32-71.0) [58] to identify protein domains from the Pfam database [59, 60] in all amino acid sequences from the representative pangenome catalogs of each species (see Pangenome analysis section). All sequences that had one or more Pfam domains were compared between datasets and two proteins were considered ‘identical’ in their domain composition when the same Pfam but no other domains were detected and the domains occurred in the same order. Proteins with identical domain compositions were clustered and visualized using the package UpsetR in R.

### Identification of CRISPR-Cas and RM systems

To identify clustered regularly interspaced short palindromic repeat (CRISPR) and their associated genes (Cas), we extracted and counted all predicted ‘cas’ and CRISPR ‘spacer’ annotations from the gff3 files generated by RAST-tk. We identified restriction-modification (RM) systems based on their Pfam domains [61], identified with Interproscan, and all coding sequences that had at least one of these RM domains were counted.

### Strain-specific gene analysis

Sequencing reads were trimmed to a quality of 20 using BBDuk (v 38.34) and mapped to each genome or MAG with a minimum identity of 0.95 using BBMap (v 38.34). The average read coverage of all samples was reduced to 100x using samtools (v 1.3.1) [62], and the MAGs with lower coverage were used with their maximum read coverages (**Tab. S7**). The identification of strain-specific genes was performed as described previously [9], for each MAG. Briefly, all genes in a genome or MAG that had lower read coverage than gammaproteobacterial marker genes were classified as strain-specific. Subsequently, all genes that had an overlap with the contig proximities (100 bp on both edges of a contig were regarded as contig proximities) were filtered out. We visualized the percentage of strain-specific genes of all coding sequences and the percentages of strain-specific genes that fall into the assembly-based core, accessory or unique genomes of a *Thioglobaceae* species using the ggplot2 package in R.

### Data availability

All MAGs and sequencing reads generated within this study were deposited at the European Nucleotide Archive under the study PRJEB36091. Accession numbers for each MAG are listed in **Tab. S1**.

## Results and Discussion

### Diversity and phylogenetic relationships within the Thioglobaceae family

To capture the genomic diversity among members of the *Thioglobaceae* family, we first collected publicly available genomes, single-cell amplified genomes (SAGs), and metagenome-assembled genomes (MAGs). We only included genomes that had > 87% completeness and < 5% contamination (see methods for details and exceptions). Altogether, this resulted in a total of 141 genomes, the most comprehensive analysis of the *Thioglobaceae* family to date (**Fig. 1, Tab. S1, S2**). These include 97 newly assembled and binned MAGs of bathymodiolin symbionts, as well as the following publicly-available genomes and MAGs: 17 MAGs and one closed genome of bathymodiolin mussel symbionts, six MAGs of sponge symbionts, three genomes of clam symbionts, one MAG of coral symbionts, and 11 MAGs, two SAG and four genomes of free-living *Thioglobaceae* lineages originating from deep-sea hydrothermal vents, cold seeps, and open water including oxygen minimum zones.

Numerous genomes and MAGs from this family were previously assigned *Candidatus* species names [63, 14, 64, 21, 41, 65]. Despite this, a family-level classification is still lacking, but is urgently needed to allow consistent naming and comparisons of species. We calculated genome phylogenies and overall genome relatedness indices (OGRI) to disentangle the taxonomic relationships between the sequenced genomes. First, to delineate species clusters we grouped all MAGs and genomes that had average nucleotide identity (ANI) of >95% (**Fig. S1**). According to these widely accepted cut-offs [42, 43], we identified 20 species clusters from the SUP05 clade and three species clusters from the sister Arctic96BD clade (**Fig. 2**). All these species clusters formed robust and divergent lineages in the phylogenomic tree constructed using 98 shared genes (**Fig 2**).

**Figure 2.**
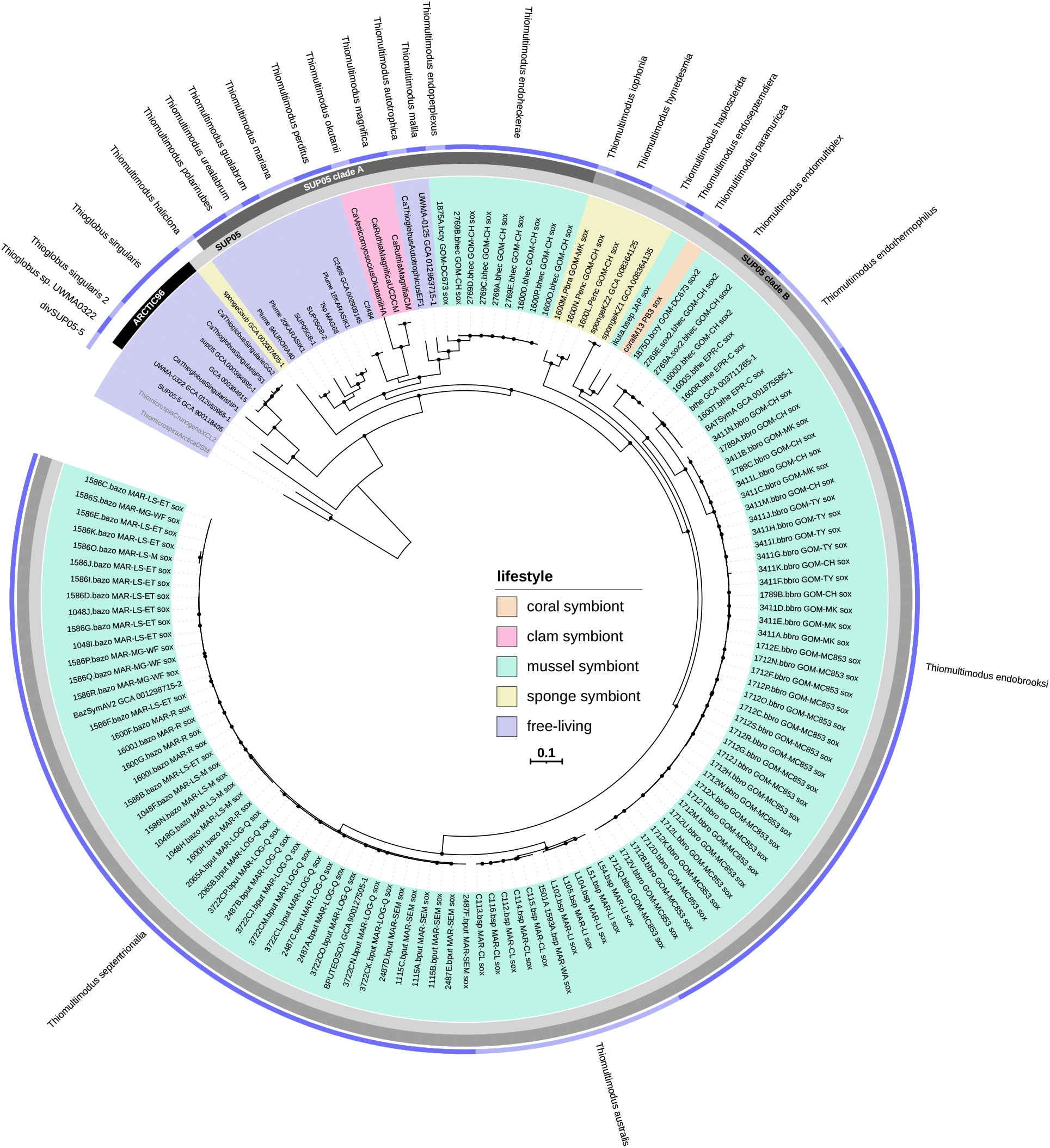
Phylogenomic tree of the *Thioglobaceae* family. The tree is based on 98 protein sequences shared among all SAGs, MAGs and genomes including the outgroup *Thiomicrospira* spp. Bootstrap support between 90-100% is represented by black circles. The inner colors in the tree represent the different lifestyles. The inner grey-colored ring indicates the two clades SUP05 and Arctic96BD. The outer grey-colored ring indicates two subclades A and B within the SUP05 clade. The outermost ring highlights the species clusters based on >95% average nucleotide identity (Fig. S1) and proposed *Candidatus* species names correspond to those described in Tab. S9.

Our results suggest that the SUP05 and Arctic96BD clades, which were both considered previously as *Ca.* Thioglobus represent two distinct genera. This is based on average amino-acid identity (AAI) and the percentage of conserved proteins (POCP), and their divergence in the phylogenomic tree. AAI of 65% has been considered a possible genus boundary cut-off [66]. As these boundaries can vary between bacterial groups [67], POCP below 50% was proposed to serve as an additional measure to differentiate bacterial genera [44]. Among the 141 genomes, we found that all members within the SUP05 clade shared AAI > 66%, whereas pairwise comparisons with Arctic96BD genomes always had AAI < 67% (**Fig. 3**). The mean of pairwise POCP value distribution between genomes of the two clades Arctic96BD and SUP05 was below 50%, whereas the mean of the distribution within the SUP05 clade was above 50%.

**Figure 3.**
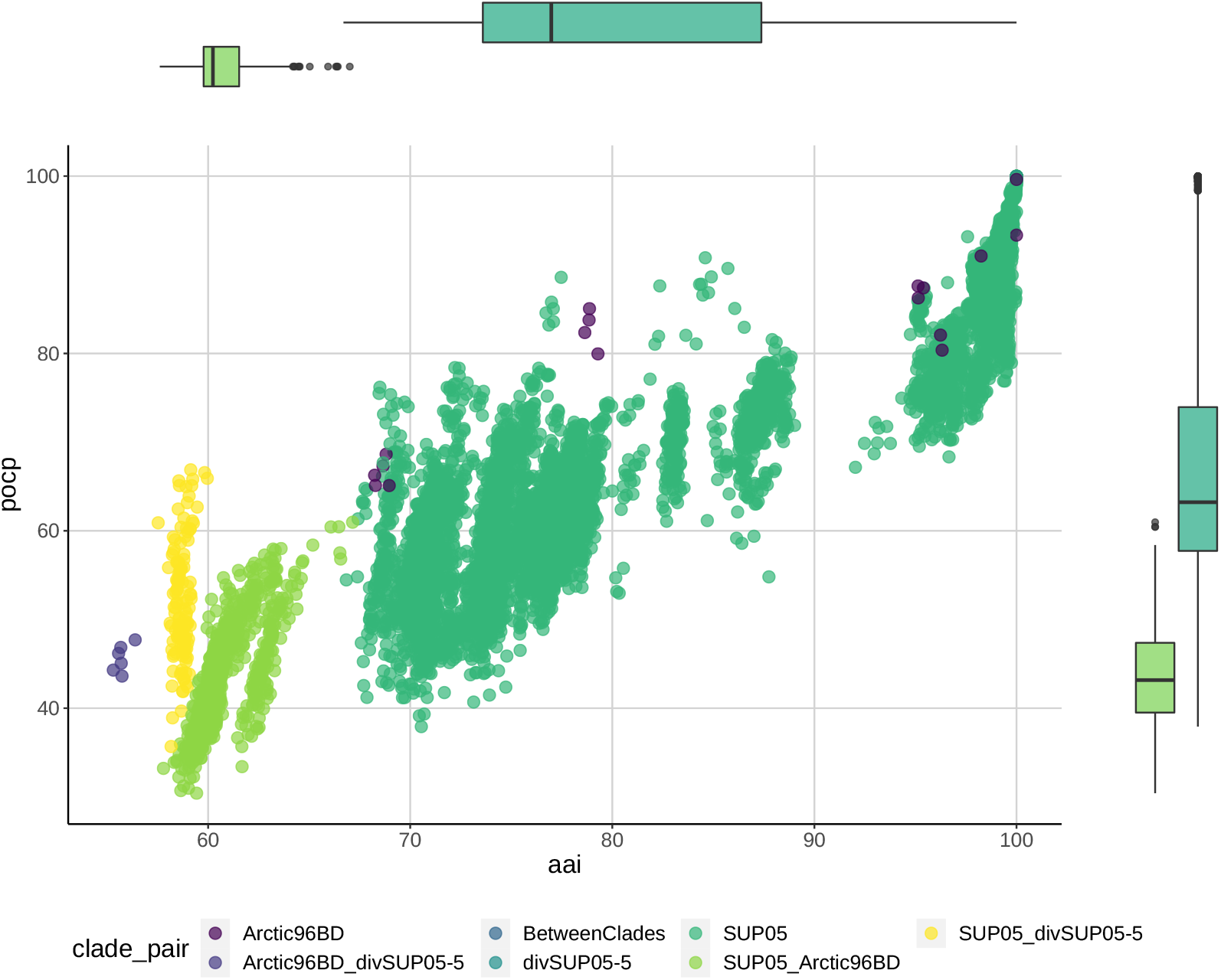
Overall genome relatedness indices among MAGs, SAGs, and genomes of the *Thioglobaceae* family. Pairwise average amino acid identity (AAI) is plotted against pairwise percentage of conserved proteins (POCP). Common genus cut-offs are 65% AAI and 50% POCP. Boxplots above and next to the scatter plot show within-clade SUP05 (dark green) and between clades SUP05 and Arctic96BD (light green) AAI and POCP, respectively.

Based on this comprehensive analysis of available genomes, using phylogeny and various OGRIs, we propose a new nomenclature for this clade including one new *Candidatus* genus and 22 new *Candidatus* species (**Fig. 2, 4;** details in **Tab. S9**). Proposing these names is important to allow the communication between scientists that study the bacteria of this family and we followed previously proposed standards for the naming of uncultivated taxa [66] (**Tab. S10**, **Tab. S4**). To establish standing species names recognized by the International Code of Nomenclature of Prokaryotes these taxa need to be further described beyond their genomic traits. One MAG was divergent to both SUP05 and Arctic96BD clades and could not be assigned to either of the two with confidence, but POCP and average amino-acid identity AAI indicated that this lineage is closer to the SUP05 clade and we refer to it as ‘divSUP05-5’ (**Fig. 3**).

**Figure 4.**
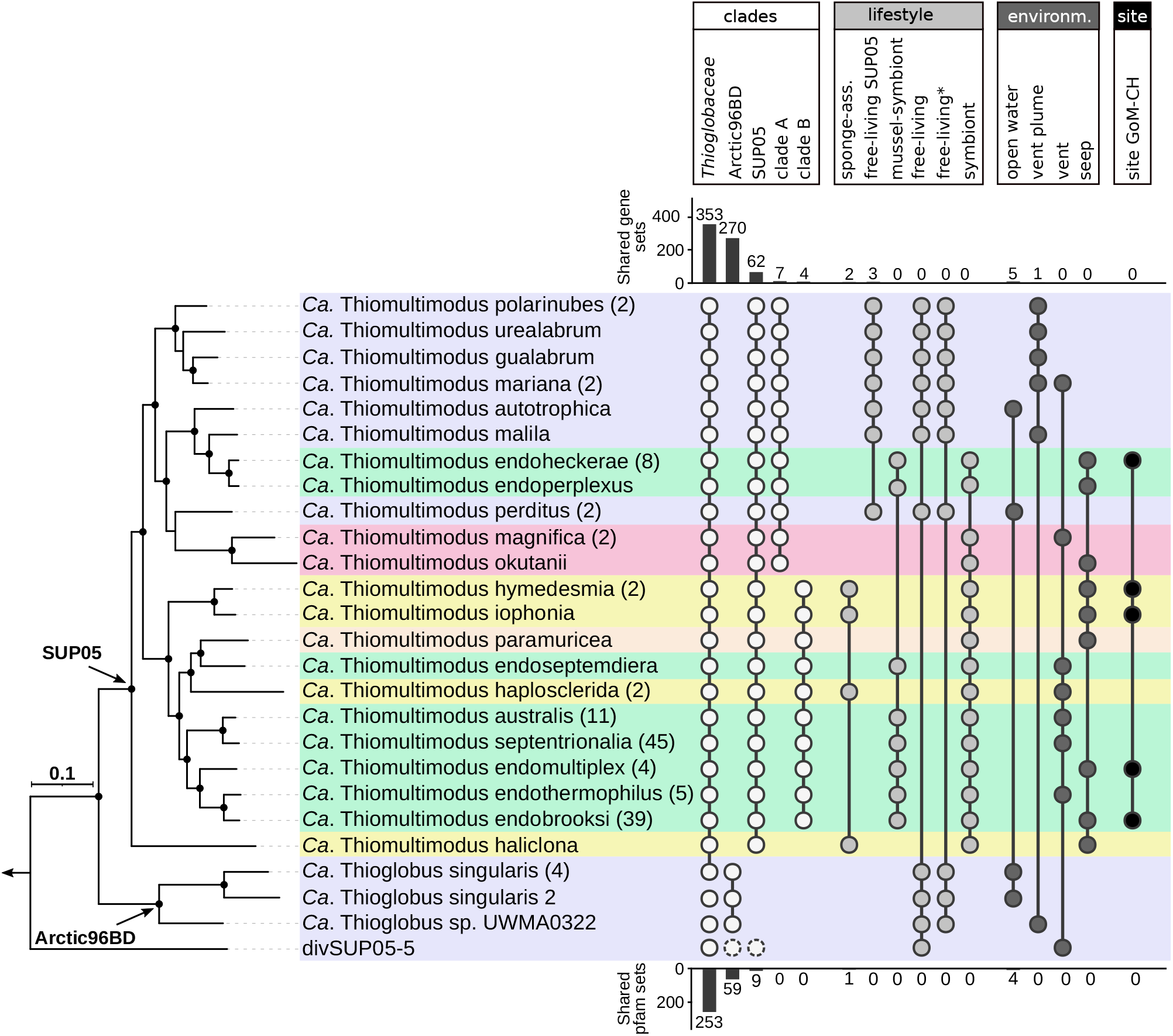
Shared gene content among groups of species within the *Thioglobaceae* family. The shared gene content was determined by comparing orthologous clustering of the species pangenome catalogues to account for potentially missing genes in single genomes (shown in bars above the plot). For shared gene sets within the SUP05- and Arctic96BD-clade, the gene content of divSUP05-5 was included in both counts as this genome could not be assigned to either of the two clades (details in Tab. S12 and S13). In addition, shared genes that had identical Pfam domain composition where identified and the numbers are shown as bars below the plot. The groups above the plots represent taxonomic clades, lifestyles, environment and site. The colors represent the lifestyles: free-living (purple), mussel-associated (green), sponge-associated (yellow), clam-associated (pink) and coral-associated (orange). The tree was collapsed at the species level and the brackets indicate the number of MAGs, SAGs, or genomes within each species.

### Metabolic differences between the Arctic96BD and SUP05 clades

We refer to the genes shared and variable among all genomes as ‘assembly-based’ core and ‘assembly-based’ accessory genomes. The assembly-based core genome of all *Thioglobaceae* species contained 353 genes. We found 62 genes unique to the SUP05 clade and 270 unique to the Arctic96BD clade (**Fig. 4, Tab. S11, S12, S13**). As indicated by the name *Thioglobaceae*, sulfur (‘*Thio*’) oxidation is considered the core metabolic feature shared by all members of this family. Surprisingly, we did not detect sulfur oxidation genes in the assembly-based core genome of the family. Although all compared *Thioglobaceae* species had the potential to oxidize sulfur, the enzymatic pathways differed substantially between the SUP05 and Arctic96BD clades. In previous cultivation experiments, sulfur oxidation and the formation of sulfur globules have been observed in *Ca.* Thioglobus singularis (Arctic96BD) and the presence of thiosulfate stimulated its heterotrophic growth [21]. However, only the adenine sulfate reductase (*aprAB*) and sulfate adenyltransferase (*sat*), which facilitate the oxidation of sulfite to sulfate, are encoded in the genome of *Ca.* Thioglobus singularis (Arctic96BD) [16]. These genomes lacked the sulfur oxidation (*sox*) and dissimilatory sulfite reduction (*dsrAB*) genes needed for the oxidation of sulfide and thiosulfate. Genes encoding the proteins that catalyze the intermediate steps of the 2-thiouridine sulfur relay were also not found. All of these genes are present in all members of the SUP05 clade and divSUP05-5 MAGs (**Tab. S12**). Instead, only the *Ca.* Thioglobus singularis lineages (Arctic96BD) encoded the proteins that are needed to use the extracellular amino acid taurine, which can be used as a sulfur source [61, 68] (**Tab. S13**). These differences in key metabolic features strengthen our conclusion that the two clades SUP05 and Arctic96BD form two distinct genera within the *Thioglobaceae* family. The Arctic96BD clade (*Ca.* Thioglobus) contains the first cultivated species *Ca.* Thioglobus singularis and we suggest retaining this genus-level naming for this clade only [21]. For the SUP05 clade, we propose the name *Ca.* Thiomultimodus gen. nov. which consists of ‘Thio’ (sulfur) and ‘multimodus’ (manifold), the latter reflecting the multiple ‘diversified’ lifestyles and hosts adopted by species belonging to this genus.

### Phylogeny is the best predictor of genetic composition within the Thioglobaceae family

The phylogenomic analysis revealed that bacterial lifestyles, such as the association with various invertebrate hosts, are intermixed across the *Thioglobaceae* tree (**Fig. 2**). Hence, we wanted to assess to what extent other factors, such as lifestyle and environment, contribute to the evolution of the family pangenome. This is of particular interest because recent data suggest that the environment is a strong driver of pangenome evolution, explaining more of the observed pangenome variation than phylogeny alone [69]. To tease apart the role of different environmental factors, we clustered the orthologous protein sequences of the ‘species pangenomes’ to determine which protein sequences are shared between species or unique to species that share a specific lifestyle or environment (**Fig. 4**). Comparing all symbiotic to the free-living lifestyles did not result in any orthologous gene set specific to either of the lifestyles. In contrast, when excluding *Ca.* Thioglobus genus (clade Arctic96BD) we detected three genes that were unique to the free-living species (**Fig. 4**). None of these genes could be reliably annotated.

We then refined the criteria used in the comparison and considered the different symbiotic lifestyles separately, as these represent three fundamentally different forms of symbioses: vesicomyid clam-associated (intracellular, vertically transmitted), bathymodiolin mussel-associated (intracellular, horizontally transmitted), and sponge-associated symbionts (unknown transmission mode and cellular location). The coral symbiont was excluded as only a single MAG was available. We identified seven orthologous genes that were exclusively shared by the vertically transmitted clam symbionts. However, these lineages form a monophyletic clade (**Fig. 2, 4**) and thus it is not possible to distinguish between phylogenetic differentiation and lifestyle-specific gene signatures. Despite having 114 MAGs available, we could not identify a single group of orthologous gene that was unique to all mussel-associated symbiont species. These data and the phylogenomic reconstruction are in line with previous conclusions suggesting that the association between *Thioglobaceae* bacteria and bathymodiolin mussels has evolved multiple times through convergent evolution [30, 27].

Intriguingly, we detected two orthologous genes specific to the three sponge symbiont species, even though they were phylogenetically quite distant from each other (**Fig. 4**). Only one gene could be properly annotated and encoded for a malate:quinone oxidoreductase (KEGG enzyme EC1.1.5.4). This enzyme converts malate into oxaloacetate and can potentially be part of various pathways including the citric acid cycle (TCA) and pyruvate metabolism. Most free-living species encoded a malate dehydrogenase (KEGG enzyme EC1.1.1.37), which can catalyze the same reaction. *Ca.* Thiomultimodus autotrophica comb. nov., *Ca*. Thiomultimodus malila sp. nov., and all clam-, and mussel-associated symbionts lacked both enzymes and thus may need to replenish intermediates into the TCA from other sources unless an uncharacterized enzyme can replace this function [70, 71]. The intracellular mussel and clam symbionts may obtain these metabolic intermediates from their hosts [70]. For the sponge symbionts the malate:quinone oxidoreductase could be an essential enzyme if they reside extracellularly in the mesohyl matrix [72], where metabolite exchange with the host may be limited. It was surprising that two of the free-living species also lacked both enzymes and the source of replenishment for the TCA intermediates in nature (or in culture) remains unclear.

We asked whether type of environment or geographic location can influence gene content in *Ca.* Thiomultimodus species. We found that the effect of the geographic location on the accessory genome is negligible, as not a single orthologous gene was specific to the sponge and mussel symbiont species that co-occur at vent site GoM-CH. We did not detect gene sets specific to the environment types hydrothermal vents (excluding plumes) or cold seeps. Genomes from lineages that occurred in vent plumes shared one orthologous gene, which could not be functionally annotated. However, we found five genes that were shared exclusively by *Ca*. Thiomultimodus and *Ca.* Thioglobus species sampled from open water environments, when excluding the divergent species divSUP05-5. Two of these genes could be annotated as ABC-type antimicrobial peptide transport system, whereas the other three genes were unknown. We conducted an additional comparison of protein domain composition to account for the fact that similar functions could be encoded by genes that are too dissimilar at the protein sequence level, and thus not detected by the orthologous clustering approach. Both approaches concurred that there were no clear differences between lifestyles and environment in the *Thioglobaceae* family (**Fig. 4**).

Finally, we inferred genome similarities based on their KEGG ortholog (KO) functional category profiles **(Fig. 5**). A distance-based redundancy analysis (dbRDA) based on Bray-Curtis dissimilarities revealed that the genomes grouped according to phylogenetic affiliation. Supporting our previous conclusions, the genera *Ca.* Thiomultimodus (SUP05 clade) and *Ca.* Thioglobus (Arctic96BD clade) were clearly separated in the ordination (**Fig. S2**). Similarly, within the *Ca.* Thiomultimodus genus MAGs clustered by their phylogeny rather than any other parameter, in agreement with the fact that genes unique to a symbiotic, mussel-associated, or free-living lifestyle were not found (**Fig. 5)**. As ‘lifestyle’ was confounded by ‘species’ we only tested the marginal effects of ‘species’ and ‘environment’ on the genomic functional profiles using permutational multivariate analysis of variance (PERMANOVA). A large proportion of variance in the dataset (31.5%) could be significantly explained by the phylogeny, whereas the factor ‘environment’ was also significant but explained only 2% of the variance (**Tab. S6C**). To visualize the effect of phylogeny on the clustering of the data, we performed an additional dbRDA for the variability introduced by the phylogenetic affiliation and showed that the clustering completely dissolved (**Fig. 5**). When conditioned for ‘environment’, the clustering in the dbRDA was diminished but not completely dissolved, showing the smaller but still significant influence of the environment compared to the effect of phylogeny (**Fig. S3**).

**Figure 5.**
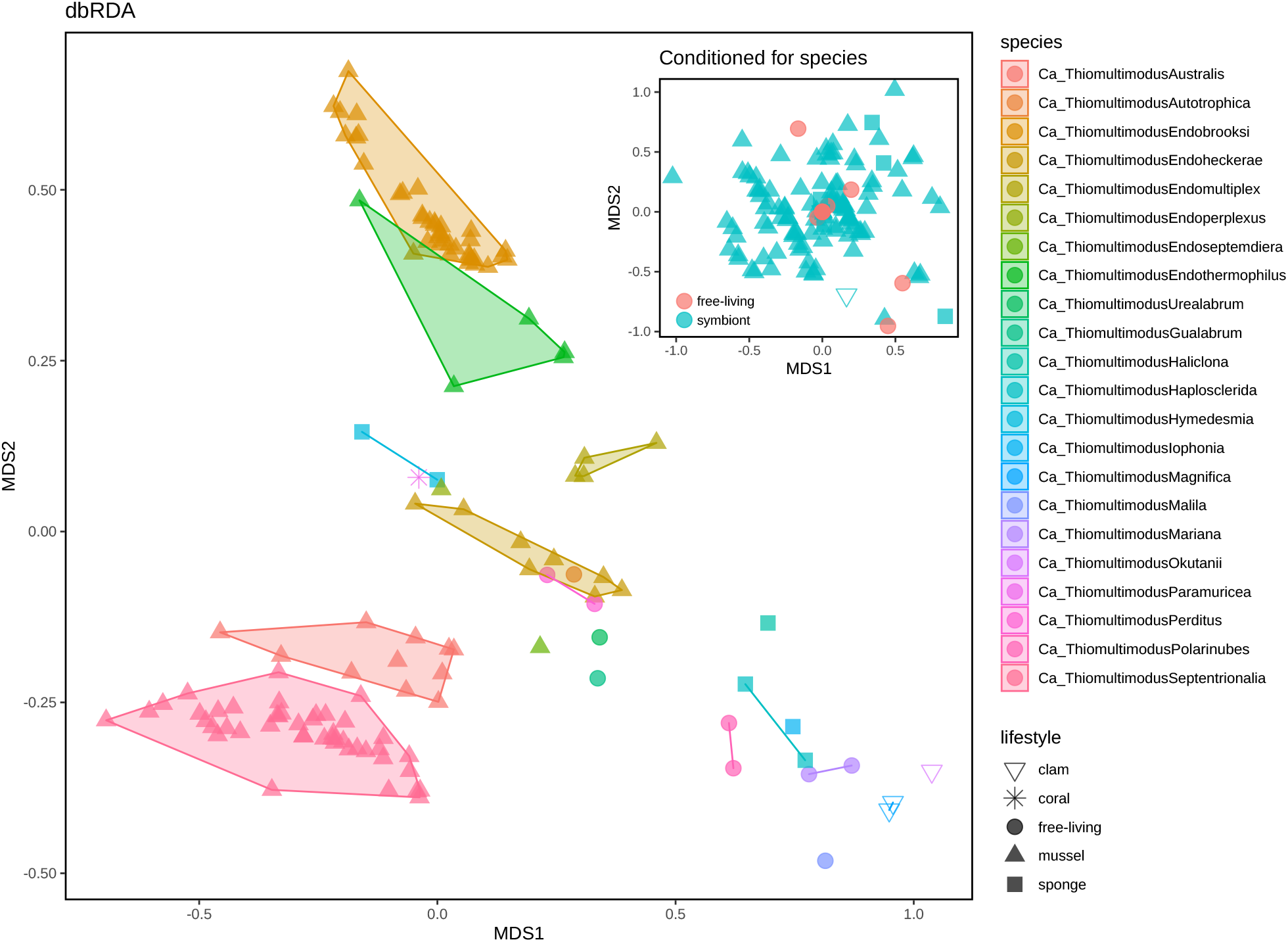
dbRDA on Bray-Curtis dissimilarities of KEGG KO profiles between MAGs and genomes within *Ca.* Thiomultimodus gen. nov. (SUP05 clade). The inset shows a dbRDA conditioned for species, hence removing the species effect on the clustering. A PERMANOVA analyses confirmed that species was the strongest significant determinant, explaining 31.5% of the clustering (Tab. S6).

Based on presence-absence analyses we observed that within the *Thioglobaceae* family, ‘lifestyle’ mostly resembled phylogenetic relatedness and appeared to be a poor predictor of overall genetic repertoires. Using the generalized odds ratio, a more permissive statistical testing approach, we then assessed whether any KEGG KO feature was enriched in bacteria with certain lifestyles. We detected 153 KOs significantly more abundant in free-living lineages and 230 KOs significantly more abundant in host-associated lineages (**Tab. S7**). In line with our previous finding, the malate dehydrogenase was enriched in the free-living species. We found restriction-modification (RM) systems, transposases, type II and type I secretion systems, phosphate-dependent regulatory system PhoB-R and the high-affinity phosphate transport system PstSCAB among 27 KOs that had the strongest difference in abundance enriched in the symbiont MAGs (LOR > 2, **Fig. 6**). Other significantly enriched KOs (0 < LOR < 2) included additional RM systems and CRISPR-Cas systems (**Tab. S7**). This data indicates that mobilome and phage-defense systems are particularly enriched in the symbiotic *Thioglobaceae*.

**Figure 6.**
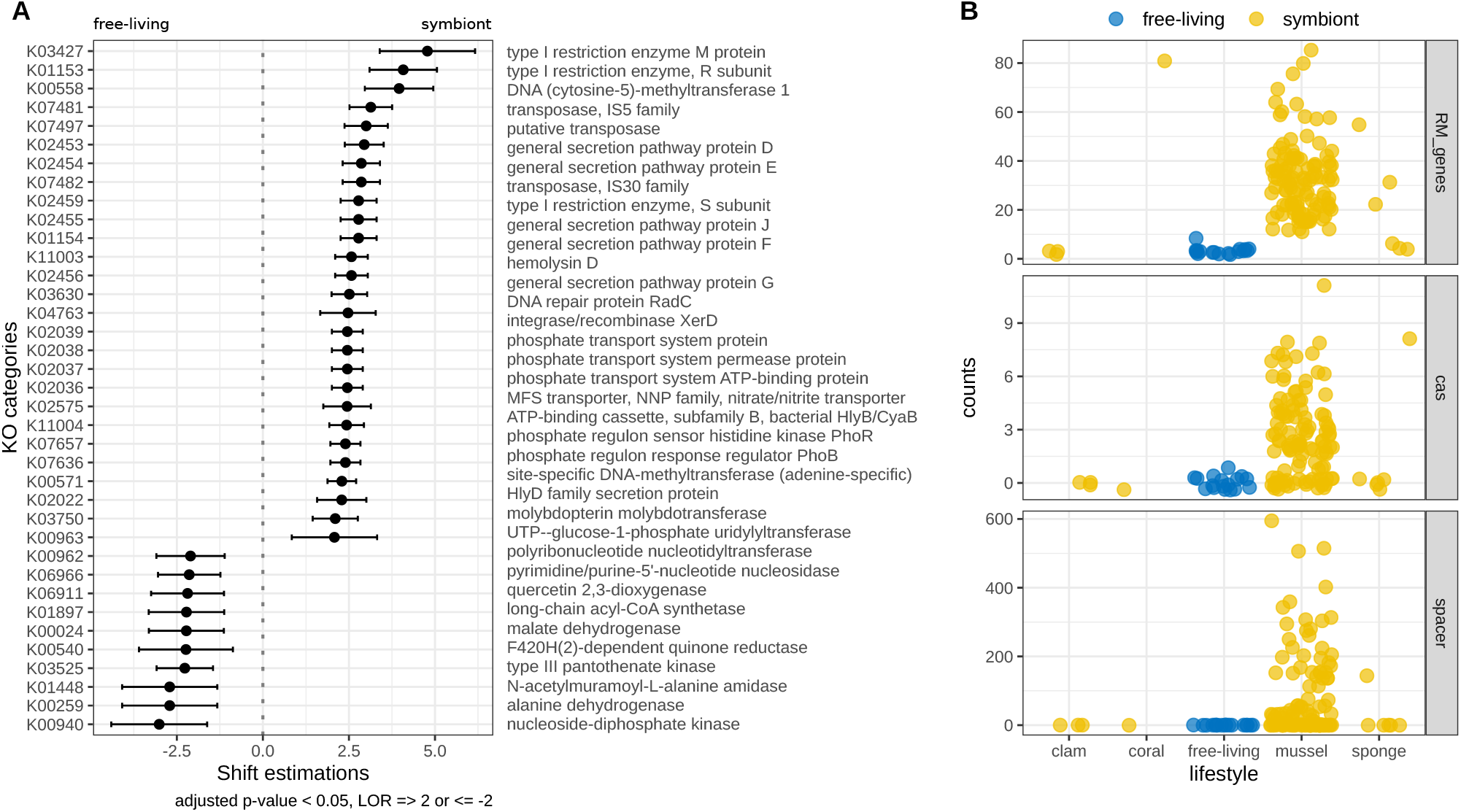
Significant differences between free-living and symbiotic *Thioglobaceae*. **A**) Log-odds ratio (LOR) and confidence intervals of KEGG KO features that were significantly different (adjusted p-value <0.05) in numbers between free-living and symbiotic lifestyles. Positive LOR indicates KOs that were enriched in symbiotic taxa and negative LOR indicates those enriched in the free-living taxa. For representative purposes, only those KOs with the strongest shift estimation (LOR > 2 or < −2) are shown (see Tab. S7 for all significant hists). **B**) Number of RM systems, CRISPR-Cas genes and CRISPR spacer sequences per genome, MAG, or SAG in the different lifestyles (symbiotic = yellow; free-living = blue).

### The hidden pangenome

Our analyses of the family pangenome revealed a considerable genomic diversity between species that is poorly explained by lifestyle and environmental factors. Considering the intra-specific variability of those species with more than 5 genomes (only the case for mussel symbionts), we observed that the assembly-based accessory genome comprised 43 to 75% of the species-pangenome, similar to the genomes of previously described medically relevant and free-living bacterial species (**Fig. S5**, [3]). This highlights that despite their intimate symbioses with a host, the accessory genomes of these species can be large. However, this assembly-based approach does not account for the strain heterogeneity potentially observed within an environmental population. This is because assemblies of short-read sequences obtained from a population formed by multiple closely-related strains are likely to result in single consensus MAGs, preventing common comparative genomic procedures to capture intra-population diversity among closely-related subspecies. Our recent study showed that such intra-population variability can be extensive and is needed to fully investigate bacterial evolution and population dynamics in a natural context [9]. In fact, accumulating evidence highlights the importance of such gene content variation for extending the adaptive potential of a population [73].

We have previously developed a read coverage-based approach that detected considerable gene content differences within natural populations of *Thioglobaceae* symbionts with identical 16S rRNA gene sequences [9]. To assess the extent of this variability across the different lifestyles within this family, we identified the genes that are present in the MAGs but are not shared among all bacterial cells within the *Ca.* Thiomultimodus genus. We found that gene content variation among co-existing strains was prevalent across lifestyles, and not specific to those living in symbiosis. Strain-specific genes comprised between 0.5 and 32% of the total protein-coding genes of genomes or MAGs (**Fig. 7**). This analysis was performed by sub-sampling the datasets to 100x average read coverage to reduce the skew due to variations in sequencing depth in different metagenomes. A few samples had read coverage below 100x (13x-97x; **Tab. S8**), therefore, the number of strain-specific genes in these metagenomes may be underestimated. As expected, most of this intra-population variable gene content was part of the assembly-based accessory genome. Surprisingly, however, a prominent number of variable genes within populations belonged to the assembly-based core genome of *Ca.* Thiomultimodus species (i.e., 3-50% for the two species with ≥ 39 available MAGs each; **Fig. 7**). This means that despite being identified in every genome assembly and thus every analyzed environmental population of a species, not every bacterial cell within those populations encoded these genes. Thus, by definition, these genes should not be regarded as part of the true core genome but instead should be regarded as accessory genomes. These findings underline the importance of extending pangenome studies to environmental populations. Only by including intra-population gene content variation can we fully capture the adaptive potential of bacterial species in nature and understand the mechanisms through which microbial genomes evolve.

**Figure 7.**
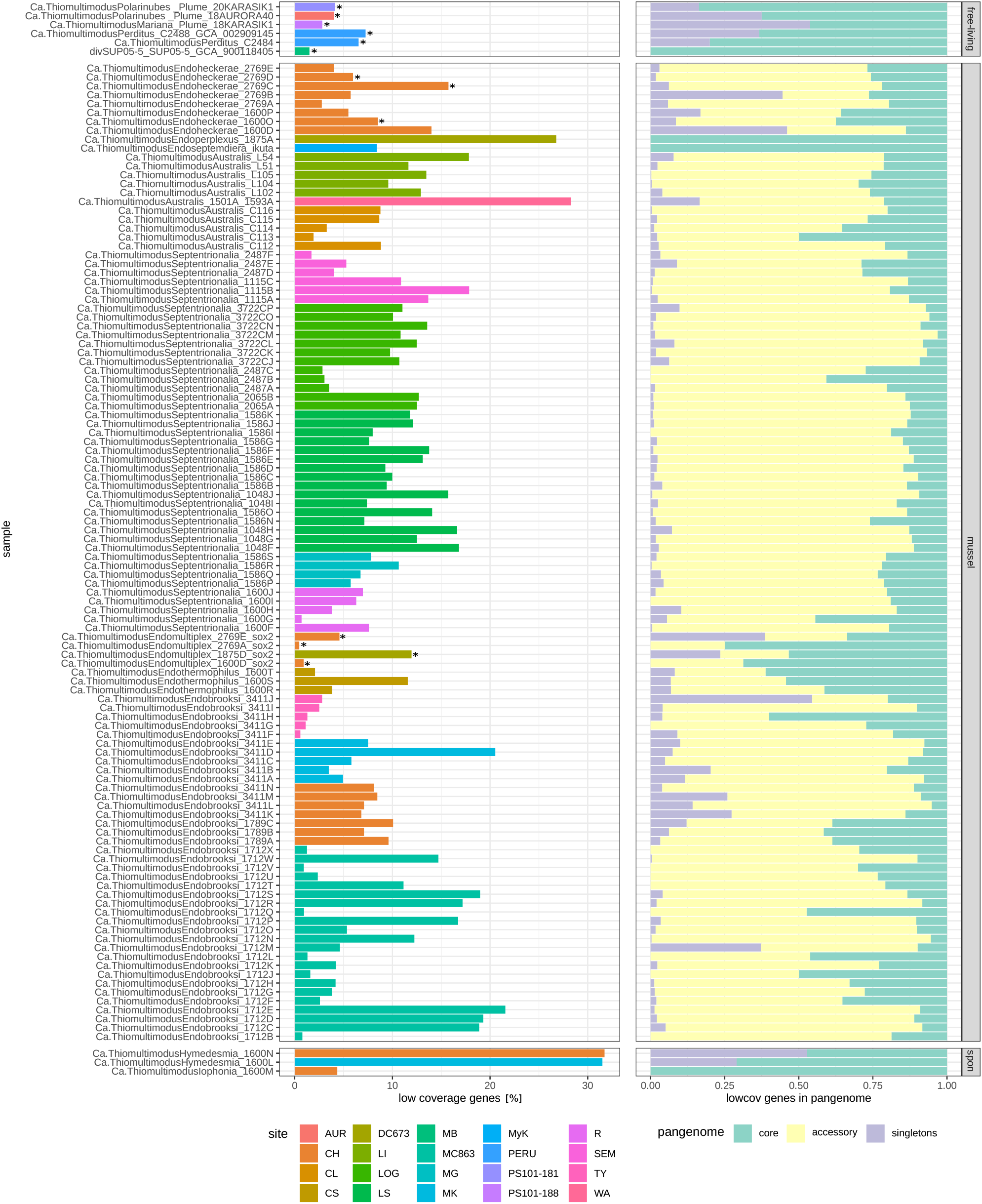
Fraction of genes [%] that were not encoded by every cell within environmental populations of *Ca.* Thiomultimodus gen. nov. species. This was retrieved from mapping the metagenomic reads to the same-sample MAGs. Most datasets were subsampled to 100x read coverage, but a few samples had lower read coverage and are marked with an asterix (precise coverage values in Tab. S8). The right panel shows what proportion of these variable genes could be assigned to the assembly-derived core genome (green), or accessory genome (yellow) and unique genes (purple) of the species. The three facets correspond to lifestyles free-living (‘free-living’), mussel-associated (‘mussel’), and sponge-associated (‘spon’).

### Genetic flexibility is key

In theoretical models, the more phylogenetically related two co-occurring strains are, the more likely they have the same growth requirements, which would often result in competition [74, 75]. We have previously suggested that genetic variation among highly related *Thioglobaceae* strains can allow them to occupy different niches and therefore co-exist [9]. Here, we show that intra-population gene content variability is pervasive within the *Thioglobaceae* family, affecting also bacterial metabolism assigned to the assembly-based core genome of some species. This highlights that common comparative genomic analyses on MAGs can miss a substantial fraction of accessory functions by falsely assigning these to the species core genome. In the *Thioglobaceae* species these included genes for processes such as nitrate respiration (e.g. *nar*, *narK*), phosphorous acquisition (e.g. *phoH*, *phnA*), iron transport (e.g. ferric iron ABC transporter) and amino acid synthesis (e.g. asparaginase, glutaminase) (**Tab. S14**). Considering evolutionary theories, some of these genes may be subject to negative frequency-dependent selection, providing a benefit only when rare in the population [76, 77]. This could apply to nitrate respiration genes that were variable in 19, out of 43 investigated, populations of a single species (*Ca.* Thiomultimodus septentrionalia), despite being assigned to the assembly-based core genome. Lower abundance of some of these genes in the population could be selected for by costs of carrying that gene, avoidance of accumulation of potentially toxic metabolic intermediates (such as nitrite during nitrogen respiration) and the exchange of intermediates between bacterial cells [78]. One explanation for detecting these variable genomic features in every MAG of a species can be that these encode essential functions for the population, while they are not essential to every single cell in the community. Therefore, these could be considered conserved core functions only at the level of the population or ‘meta-organism’, the latter representing the association between a host and its symbionts [79], and not the core genome of a species.

The extensive intra-specific genomic variability we detected within the *Thioglobaceae* family aligns with the evolvability concept, which is considered as the potential of a population to evolve adaptive solutions to unknown future conditions [80]. It is debated in the theoretical literature if and how evolvability itself can be favored by selection [80, 81]. Theory would predict that if the cost of encoding a gene in a genome is greater than the benefits gained, it will tend to be lost [82, 83]. Genes for which the cost-benefit ratio hovers around zero, providing slight total benefits or slight total costs, may be ‘kept’ available at the population level, as they might provide a potential adaptive benefit when the environment changes. The larger the gene pool, particularly when rampant horizontal gene transfer occurs (see below), the higher the chances that the population encodes a metabolism that confers a stronger fitness benefit under new conditions. In this way, these genes can either be passed on and acquired by a greater proportion of the population, or the cells encoding them can proliferate more successfully than those without. It is worth mentioning that within this scenario some of the variable genes we observed within a population may never provide a selective advantage.

Such heterogeneity in a population may be widespread in natural environments that are heterogeneous and change over time, whereas homogeneous environments likely lead to the purging of genomic heterogeneity by purifying selection [83]. Evolvability therefore depends on two factors: genomic innovation and ecological opportunity. Bacterial species belonging to *Ca.* Thiomultimodus often thrive in heterogeneous environments with steep chemical gradients and rapid change, such as the oxygen minimum zones where dynamic water currents and mixing expose free-living populations to fluctuating conditions. This can also be the case for the horizontally transmitted mussel symbionts at hydrothermal vents where extreme changes in conditions may result from environmental fluctuations and exposure to within-host and outside-host environments. The key to *Ca.* Thiomultimodus’ success may be the ability to maintain enormous genomic flexibility, from shorter ecological time scales with mosaics of strains occupying distinct micro-niches within a population, to evolutionary time scales resulting in genus-wide plasticity in gene content [16]. *Ca.* Thiomultimodus bacteria partition these capabilities into distinct strains with small genomes (**Fig. S5**) rather than one versatile strain that carries the costly potential to deal with all possible changing conditions.

One factor that can increase genomic plasticity and possibly lead to convergent evolution of similar lifestyles with different genetic setups, is horizontal gene transfer (HGT). Metabolic genes that could enable niche-partitioning, such as the hydrogenase, are affected by HGT in symbiotic *Ca.* Thiomultimodus gen. nov (SUP05) bacteria [71]. This, as well as the duplication and loss of genes, can lead to variation in gene content and increase the accessory genome of bacterial species.

### Increased exchange of genetic material in mussel and sponge symbionts

Mechanisms that increase rates of genetic innovation and exchange would increase the evolvability of a species. One example of such mechanisms is phages-mediated HGT and in fact, massive phage infection has been shown for free-living *Ca.* Thiomultimodus bacteria where key sulfur-oxidation genes are found in the phage gene pool [84, 85, 23]. Thus, phages likely are among the driving forces of genetic exchange in this genus, possibly contributing to their evolvability [90]. In agreement with this, RM and CRISPR-Cas systems were relatively enriched in mussel, sponge and coral symbionts compared to the free-living lineages (**Fig. 6).** CRISPR-Cas systems are prokaryotic defense mechanisms against bacteriophages and foreign DNA [86]. Except for *Ca.* Thiomultimodus endoheckerae sp. nov., we found *cas* genes in all mussel symbiont species, with high variability in numbers between genomes and MAGs. Such intra-specific variability in phage defense mechanisms is commonly observed in bacterial species leading to a flexible pan-immune system within populations [87]. Two of the sponge-associated species also encoded *cas* genes. Except for a single *cas* gene in the divergent free-living lineage divSUP05-5, neither the free-living nor the coral and clam symbionts in the T*hioglobaceae* encoded any *cas* gene. Similarly, RM systems that have been described to be involved in phage defense, increased genomic variation, control of HGT and stabilization of genomic islands [88], were more abundant in mussel and sponge symbionts than in clam symbionts and free-living lineages. This is in line with previous observations [89, 90, 32, 91] and strongly suggests an important role of these mechanisms for the evolution of a facultative host-associated lifestyle. HGT in mussel, sponge and coral symbionts appears rampant even though the host may offer a sheltered environment from phage predation. Possibly, the free-living stage during horizontal transmission may provide phages a window of opportunity to infect bacterial cells that will later, inside a host, become part of a high-density population. Alternatively, phages might reach bacteria that reside intracellularly in eukaryotic cells [92]. The lack of CRISPR-Cas systems in the clam symbionts is likely attributed to the vertical transmission which is accompanied by bottlenecks and subsequent genome reduction, which, possibly led to a loss of these defense systems by genetic drift [93].

A higher number of RM and CRISPR-Cas is generally subjected to and associated with a higher rate of HGT [87, 94]. For example, naturally competent bacteria with small genomes, including *Helicobacter pylori*, have been characterized by increased numbers of RM systems [88, 94]. This suggests that mussel and sponge symbionts experienced an increased rate of HGT, although it is unclear whether RM and CRISPR-Cas systems have an active role in HGT or are merely a consequence. In addition to the uptake of foreign DNA, both systems have also been shown to increase heterogeneity within bacterial populations through self-targeting affecting gene expression [95, 88]. Association with a host could potentially increase the selective pressure towards increased rates of HGT and population heterogeneity, adding the ‘host-associated lifestyle’ as an additional level of selection, which has to be separated from the idea that symbiont and host evolve only as a ‘unit’ which is very much debated in the field [96]. Such an additional layer of selection might exist because, first, a host at hydrothermal vents and seeps harboring a more flexible and genomically adaptable symbiont population might be more successful as has been suggested for this and other systems before [9, 97]. And second, horizontally transmitted symbionts (such as the mussel symbionts) experience extreme changes in conditions between a within-host and outside-host environment were population heterogeneity of distinct genetic features may be an advantage to the symbionts. Compared to a permanently free-living bacterial population, these two factors could increase selective pressure towards mechanisms that increase genomic heterogeneity and thus evolvability within the symbiont population.

## Conclusion

The *Thioglobaceae* family (commonly referred to as SUP05 and Arctic96BD clade) showed a fine-scale genomic diversity that is common among all its members and was pervasive throughout all taxonomic ranks and lifestyles investigated. We discovered a ‘hidden’ intra-population diversity in symbiotic and free-living lifestyles throughout the novel genus *Ca.* Thiomultimodus, that is completely overlooked when standard comparative genomics approaches are used. The bacterial variants with different gene content in these populations can be considered ‘puzzle pieces’ that resemble a composition of ‘core functions’ as a consortium instead of one single versatile strain. We propose that selection for high evolvability by enhancing intra-population heterogeneity paired with a fluctuating or variable environment is the key to the global success of this genus. Our data indicate that population heterogeneity might be more advantageous in some of the host-associated lifestyles than free-living ones and can be explained only if increased HGT and heterogeneity are selected for at the level of the host-association. In agreement with our findings, other widespread marine species have recently been shown to have intra-population variation at the nucleotide level underlining the fact that such sub-specific variability may be a ubiquitous phenomenon across bacterial groups and lifestyles in nature [4]. Our findings of intra-population variation in gene content, the ‘hidden pangenome’, show that it is not only possible but necessary to study population pangenomes of cultivated and uncultivated bacteria in their natural environment. These variable and low-frequency traits are the foundation for the adaptability of free-living and host-associated bacteria.

## Acknowledgements

We thank the captains, crews and ROV teams on the cruises BioBaz (2013), ODEMAR (2014), M126 (2016), M78-2 (2009), NA58 (2015), NA43 (2014), M114-2 (2015), AT26-23 (2014), ATA57 (2008) and M93 (2013) on board of the research vessels Pourquoi Pas?, FS Meteor, E/V Nautilus, and L’Atalante and the chief scientists on these research expeditions. This study was funded by the Max Planck Society, the MARUM DFG-Research Center / Excellence Cluster “The Ocean in the Earth System” at the University of Bremen, the German Research Foundation, an ERC Advanced Grant (BathyBiome, 340535), and a Gordon and Betty Moore Foundation Marine Microbial Initiative Investigator Award to ND (Grant GBMF3811).

