## SupplementaryMaterial for "The hidden pangenome: comparative genomics reveals pervasive diversity in symbiotic and free-living sulfur-oxidizing bacteria": Ansorge_etal_supplement.pdf

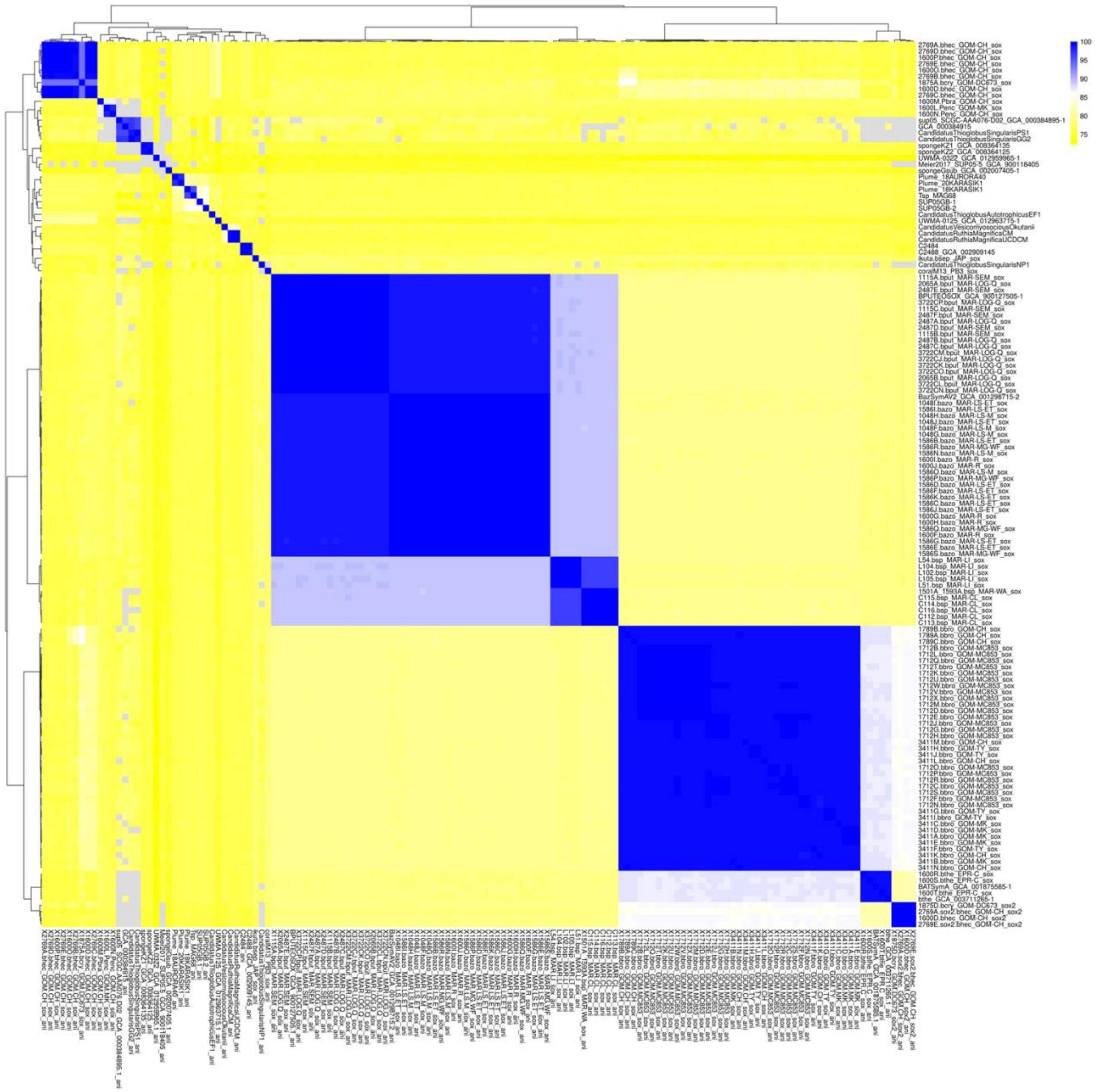


**Figure 1| Pairwise average nucleotide identities (ANI) between MAGs and genomes within the *Thioglobaceae* family.** ANI of <95% was considered the species cutoff, resulting in 26 distinct species clusters in the family.

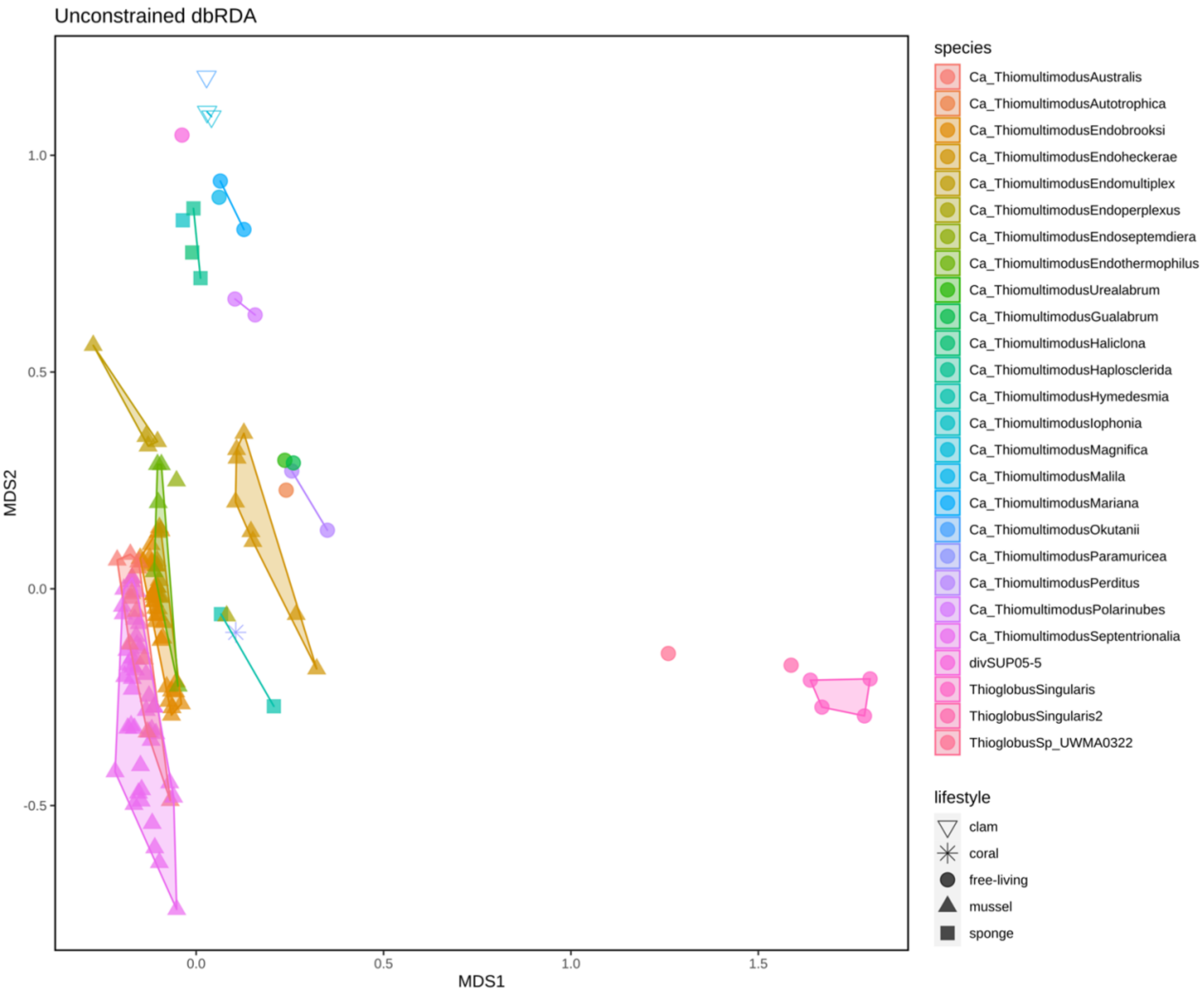


**Figure 2 dbRDA on Bray-Curtis dissimilarities of KEGG KO profiles between MAGs and genomes within the *Thioglobaceae* family.**

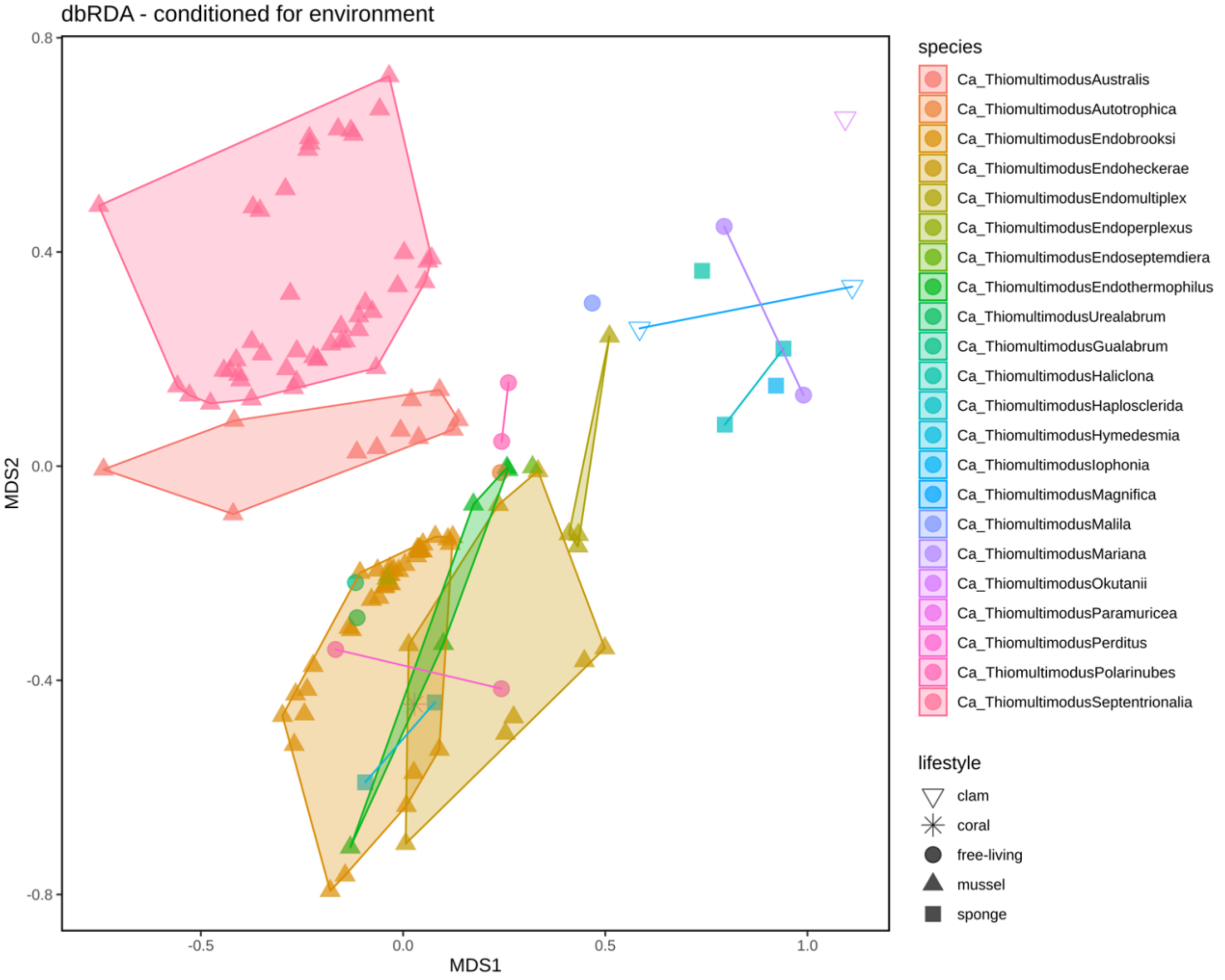


**Figure 3 | dbRDA on Bray-Curtis dissimilarities of KEGG KO profiles between MAGs and genomes within *Ca.* Thiomultimodus gen. nov. (SUP05 clade).** The dbRDA was conditioned for environment, hence removing the environment effect on the clustering. A PERMANOVA analyses confirmed that environment has a significant effect on the clustering (Tab. 6).

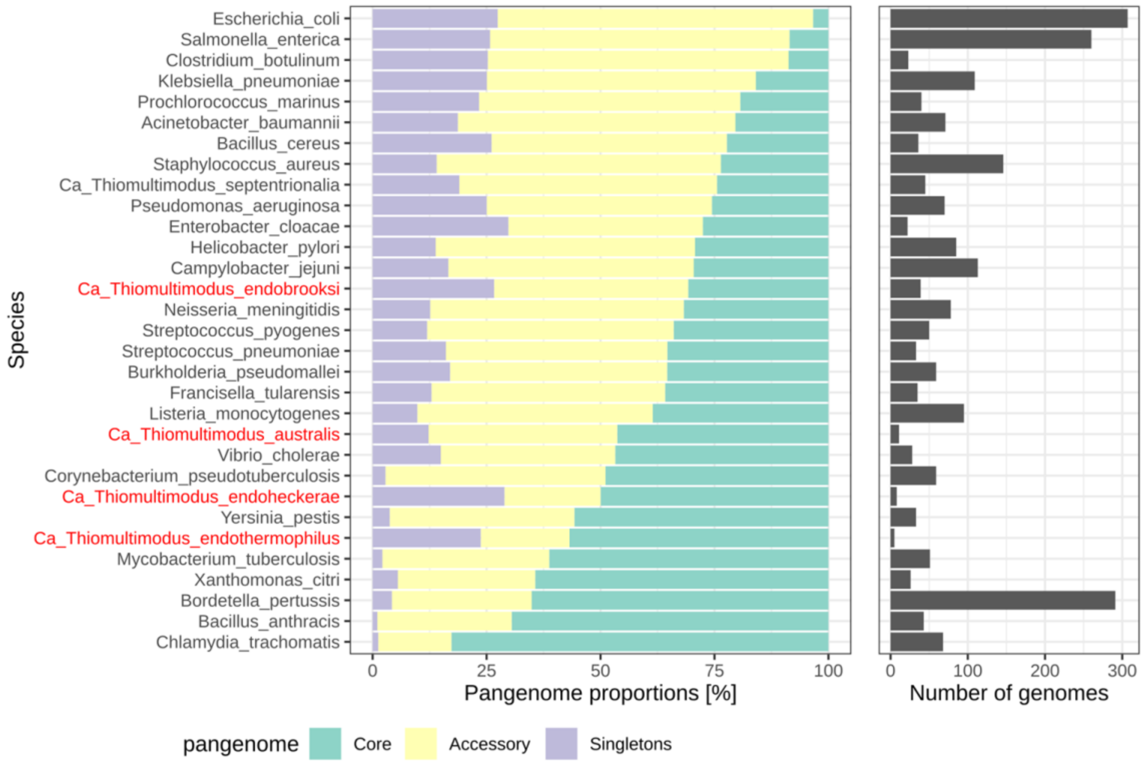


**Figure 4 | Assembly-derived pangenome of *Ca.* Thiomultimodus gen. nov. (SUP05) species compared to known species pangenomes.** Only species with 5 or more MAGs were included and the number of genomes available per species is shown in the right panel. *Ca.* Thiomultimodus spp. are highlighted in red. *obtained from (Ding, Baumdicker, and Neher 2018)

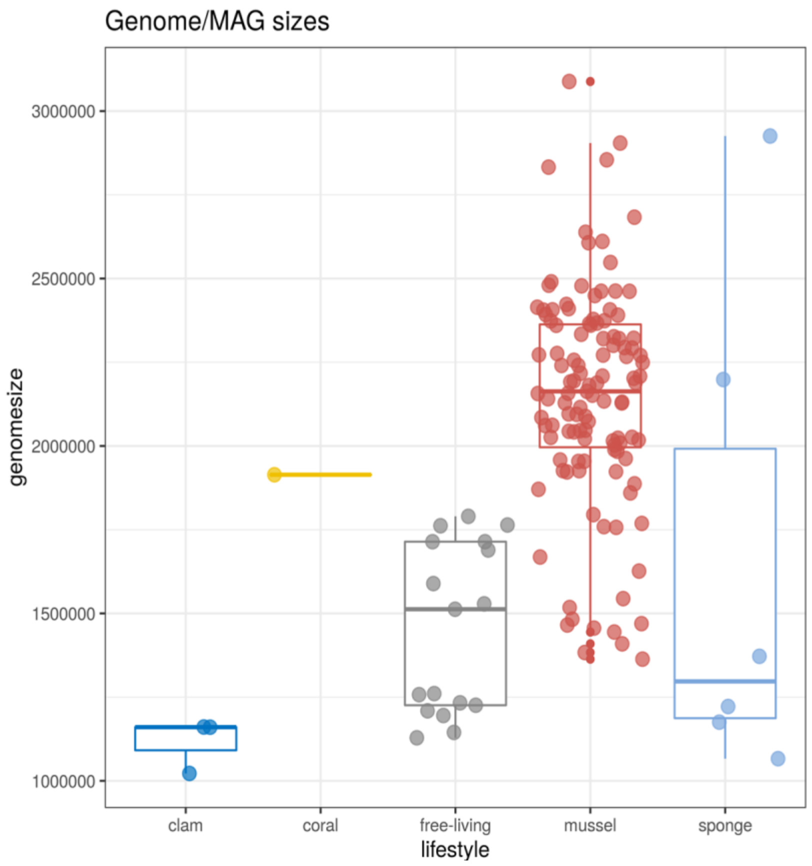


**Figure 5 | Genome sizes [bp] of genomes and MAGs in *Thioglobaceae* family.**
