## SupplementaryMaterial for "The hidden pangenome: comparative genomics reveals pervasive diversity in symbiotic and free-living sulfur-oxidizing bacteria": Tab_S9.docx

**Table S9 | Proposed *Candidatus* (*Ca.*) genus and *Ca.* species names of mostly uncultivated* taxa in SUP05 clade.** Some proposed Ca. species names will be newly generated and others will be modified from previously proposed names to conform with the proposed novel *Ca.* genus.

| **Proposed genus name** | **Description of genus name** |
| --- | --- |
| *Ca.* Thiomultimodus gen. nov. | We are proposing a new genus name based on the results in this study. This new genus encompasses the former SUP05 clade but excludes the Arctic96BD clade. Formerly cultivated species of both clades have been named *Ca.* Thioglobus spp. We are proposing to split this group into the genus *Ca.* Thioglobus (Arctic96BD clade) and the novel genus *Ca*. Thiomultimodus gen. nov., named after ‘Thio’ referring to the main energy source of reduced sulfur compounds; and ‘multimodus’ which means ‘manifold’ and refers to the different lifestyles within this genus.  Etymology:  ‘thios’ (Gr. noun): sulfur  ‘multimodus’ (L. masc. adj.): manifold |

| **Previously** | **Proposed species name** | **Description of species name** |
| --- | --- | --- |
|  | *Ca.* Thiomultimodus septentrionalia sp. nov. | This species of mussel symbionts of the novel genus *Ca.* Thiomultimodus gen. nov. was named ‘septentrionalia’ (latin for ‘north’) as this symbiont species colonized multiple *Bathymodiolus* species from the northern Mid Atlantic Ridge  Etymology:  ‘septentrionalia’ (L. n. noun): multiple regions in the north (pl.) |
|  | *Ca.* Thiomultimodus australis sp. nov. | This species of mussel symbionts of the novel genus *Ca.* Thiomultimodus gen. nov. was named ‘australis’ (latin for ‘southern’) as this symbiont species colonized multiple *Bathymodiolus* species from the southern Mid Atlantic Ridge  Etymology:  ‘australis’ (L. masc. adj): southern |
|  | *Ca.* Thiomultimodus endobrooksi sp. nov. | This species of the novel genus *Ca.* Thiomultimodus gen. nov. was named after the host species (*Bathymodiolus brooksi*) with the prefix ‘endo’ to represent ‘endosymbiont’.  Etymology:  ‘endon’ (Gr. noun): within  ‘brooksi’: after host species *Bathymodiolus brooksi* |
|  | *Ca.* Thiomultimodus endoheckerae sp. nov. | This species of mussel symbionts of the novel genus *Ca.* Thiomultimodus gen. nov. was named after the host species (*Bathymodiolus heckerae*) with the prefix ‘endo’ to represent ‘endosymbiont’.  Etymology:  ‘endon’ (Gr. noun): within  ‘heckerae’: after host species *Bathymodiolus heckerae* |
|  | *Ca.* Thiomultimodus endoperplexus sp. nov. | This species of mussel symbionts of the novel genus *Ca.* Thiomultimodus gen. nov. was named ‘perplexus’ (latin for ‘cryptic’) as this symbiont species colonized a so far undescribed, cryptic *Bathymodiolus* species from the Gulf of Mexico, with the prefix ‘endo’ to represent ‘endosymbiont’.  Etymology:  ‘endon’ (Gr. noun): within  ‘perplexus’ (L. masc. adj.): cryptic |
|  | *Ca.* Thiomultimodus endothermophilus sp. nov. | This species of mussel symbionts of the novel genus *Ca.* Thiomultimodus gen. nov was named after the host species (*Bathymodiolus thermophilus*) with the prefix ‘endo’ to represent ‘endosymbiont’.  Etymology:  ‘endon’ (Gr. noun): within  ‘thermophilus’: after host species *Bathymodiolus thermophilus* |
| *Ca.* Thiodubiliella endoseptemdiera | *Ca.* Thiomultimodus endoseptemdiera comb. nov. | This species of mussel symbionts has been proposed as *Ca.* Thiodubiliella endoseptemdiera (Russel et al., 2020). Since we define a new and different genus in our study, we combined the same species name with the herein novel genus *Ca.* Thiomultimodus gen. nov. |
|  | *Ca.* Thiomultimodus endomultiplex sp. nov. | This species of mussel symbionts of the novel genus *Ca.* Thiomultimodus gen. nov. was named ‘multiplex’ (latin for ‘complex’) as this symbiont species colonized multiple different *Bathymodiolus* species from the Gulf of Mexico. This symbiont also reside alongside other *Ca.* Thiomultimodus spp. symbionts in the same host individual and species. Both facts were reasons for the choice of name.  Etymology:  ‘endon’ (Gr. noun): within  ‘multiplex’ (L. masc. adj.): complex, having many layers |
|  | *Ca.* Thiomultimodus urealabrum sp. nov. | This free-living species of the novel genus *Ca.* Thiomultimodus gen. nov. was named after the sampling location Guaymas ‘Basin’ with the suffix ‘urea’ referring to the gene operon encoding a urease (Antharaman et al.; 2013). It corresponds to the ‘GB-1’ bin described in the mentioned publication.  Etymology:  ‘urea’: refers to the urease operon that encodes a urease which can break down urea  ‘labrum’ (L. m. noun.): large basin |
| GB-2 | *Ca.* Thiomultimodus gualabrum sp. nov. | This free-living species of the novel genus *Ca.* Thiomultimodus gen. nov. was named after the sampling location ‘Guaymas’ basin with the suffix ‘labrum’ (latin for ‘large basin’). It corresponds to the ‘GB-2’ bin described in Antharaman et al. (2013).  Etymology:  ‘gua’: refers to ‘*Gua*ymas Basin’  ‘labrum’ (L. m. noun): large basin |
|  | *Ca.* Thiomultimodus polarinubes sp. nov. | This free-living species of the novel genus *Ca.* Thiomultimodus gen. nov. was named as a combination of ‘polaris’ (latin for ‘polar’) and ‘nubes’ (latin for ‘cloud’) as this species was sampled at hydrothermal plumes (cloud analogy) at northern sites in polar waters.  Etymology:  ‘polaris’ (L. m. adj.): polar  ‘nubes’ (L. fem. noun): cloud |
|  | *Ca.* Thiomultimodus mariana sp. nov. | This free-living species of the novel genus *Ca.* Thiomultimodus gen. nov. was named after the origin of one of the MAGs from hydrothermal venting fluids at the ‘Mariana’ back-arc spreading center.  Etymology:  ‘mariana’: refers to ‘*Mariana*’ back-arc spreading center |
| *^U^*Thioglobus perditus | *Ca.* Thiomultimodus perditus comb. nov. | This free-living species has been proposed as *^U^*Thioglobus perditus (Callbeck et al, 2018). Since we define a new and different genus in our study, we combined the same species name with the herein novel genus *Ca.* Thiomultimodus gen. nov. |
| *Thioglobus autotrophica* | *Ca.* Thiomultimodus autotrophica comb. nov. | This free-living species has been proposed as *Ca.* Thioglobus autotrophica (Shah and Morris, 2015). Since we define a new and different genus in our study, we combined the same species name with the herein novel genus *Ca.* Thiomultimodus gen. nov. |
|  | *Ca.* Thiomultimodus hymedesmia sp. nov. | This species of sponge symbionts of the novel genus *Ca.* Thiomultimodus gen. nov. was named after the sponge host *Hymedesmia methanophila*  Etymology:  ‘hymedesmia’: after host species *Hymedesmia methanophila* |
|  | *Ca.* Thiomultimodus iophonia  sp. nov. | This species of sponge symbionts of the novel genus *Ca.* Thiomultimodus gen. nov. was named after the sponge host *Iophon methanophila*  Etymology:  ‘iophonia’: after host species *Iophon methanophila* |
|  | *Ca.* Thiomultimodus haplosclerida sp. nov. | This species of sponge symbionts of the novel genus *Ca.* Thiomultimodus gen. nov. was named after the sponge host *Haplosclerida* sp.  Etymology:  ‘*haplosclerida*’: after host species *Haplosclerida* sp. |
|  | *Ca.* Thiomultimodus paramuricea sp. nov. | This species of coral symbionts of the novel genus *Ca.* Thiomultimodus gen. nov. was named after the coral host *Paramuricea* sp.  Etymology:  ‘paramuricea’: refers to coral host *Paramuricea* sp. |
| *Ca.* Ruthia magnifica | *Ca.* Thiomultimodus magnifica comb. nov. | This species of clam symbionts has been proposed as *Ca.* Ruthia magnifica (Newton et al., 2007). Since we define a new and different genus in our study, we combined the same species name with the herein novel genus *Ca.* Thiomultimodus gen. nov. |
| *Ca*. Vesicomyosocius okutanii | *Ca.* Thiomultimodus okutanii comb. nov. | This species of clam symbionts has been proposed as *Ca.* Vesicomyosocius okutanii (Kuwahara et al., 2008). Since we define a new and different genus in our study, we combined the same species name with the herein novel genus *Ca.* Thiomultimodus gen. nov. |
|  | *Ca.* Thiomultimodus haliclona sp. nov. | This species of sponge symbionts of the novel genus *Ca.* Thiomultimodus gen. nov was named after the sponge host *Haliclona cymaeformis* |
|  | *Ca.* Thiomultimodus malila sp. nov. | This free-living species of the novel genus *Ca.* Thiomultimodus gen. nov was named after the sampling location ‘Tui Malila’ hydrothermal plume Etymology:  ‘malila’: refers to ‘Tui *Malila*’ |

*only *Ca.* Thiomultimodus autotrophica comb. nov. has been cultured
